# Excessive Mechanotransduction in Sensory Neurons Causes Joint Contractures in a Mouse Model of Arthrogryposis

**DOI:** 10.1101/2022.06.07.495164

**Authors:** Shang Ma, Adrienne E. Dubin, Luis O. Romero, Meaghan Loud, Alexandra Salazar, Yu Wang, Alex T. Chesler, Katherine Wilkinson, Valeria Vásquez, Kara L. Marshall, Ardem Patapoutian

**Affiliations:** Howard Hughes Medical Institute, Department of Neuroscience, Dorris Neuroscience Center, Scripps Research, La Jolla, CA 92037, USA; Department of Physiology, College of Medicine, University of Tennessee Health Science Center, Memphis, TN, USA; Integrated Biomedical Sciences Graduate Program, College of Graduate Health Sciences, University of Tennessee Health Science Center, Memphis, TN, USA; National Institute of Neurological Disorders and Stroke, National Institutes of Health, Bethesda, MD, USA; National Center for Complementary and Integrative Health, National Institutes of Health, Bethesda, MD, USA; Department of Biological Sciences, San Jose State University, San Jose, CA, USA; Department of Neuroscience, Baylor College of Medicine, Houston, TX, USA

## Abstract

Distal arthrogryposis (DA) is a collection of rare disorders characterized by congenital joint contractures. Most DA mutations are in muscle- and joint-related genes, and the anatomical defects originate cell-autonomously within the musculoskeletal system. However, gain-of-function (GOF) mutations in PIEZO2, a principal mechanosensor in somatosensation, cause DA subtype 5 via unknown mechanisms. We show that expression of a GOF PIEZO2 mutation in proprioceptive sensory neurons mainly innervating muscle spindles and tendons is sufficient to induce DA5-like phenotypes in mice. Overactive PIEZO2 causes anatomical defects via increased activity within the peripheral nervous system during postnatal development. Remarkably, Botox and a dietary fatty acid that modulates PIEZO2 activity markedly reduce DA5-like deficits. This reveals an unexpected role for somatosensory neurons: excessive mechanosensation within these neurons disrupts musculoskeletal development.

## Main text

Distal Arthrogryposis (DA) is a rare disorder with congenital contractures primarily affecting joints. It is estimated to afflict ∼1 in 3,000 individuals worldwide and usually requires invasive surgeries to alleviate the symptoms (1). The etiology of DA or the related arthrogryposis multiple congenita (AMC) may involve environmental factors such as decreased intrauterine movement (2). It also has strong genetic roots and various mutations have been identified in DA patients. These mutations occur in genes important for musculoskeletal function such as those in myosin heavy chain gene family (3). DA5 is autosomal dominant with distinct clinical features including ophthalmoplegia, and in some cases restrictive lung disease, in addition to contractures in distal joints (4). Gain-of-function (GOF) mutations in PIEZO2, a mechanically activated ion channel, have been found in DA5 patients (5). PIEZO2 is the principal mechanosensor in somatosensory neurons for detecting pressure and translating this stimulus into neuronal signals. It underlies touch sensation (6) and proprioception (the sense of where one’s limbs are in space) (7), as well as mechanosensory processes in internal organs such as lung, aorta, and bladder (8–10). Identification of *PIEZO2* mutations in DA5 patients was surprising, as a direct role for this ion channel in muscles or tendons has not been described. We set out to ask how PIEZO2 might contribute to these DA5 phenotypes.

We engineered mice that can conditionally express a human-equivalent GOF *Piezo2* mutant to study the disease mechanism for DA5. One such GOF mutation is in the C-terminus of PIEZO2 protein and causes significantly slower channel inactivation (larger inactivation time constant, tau), resulting in more ions conducted compared to wild type in response to a given mechanical stimulus (5). We designed a knock-in strategy that allows Cre recombinase-dependent replacement of a wild type C-terminal exon of PIEZO2 with the same exon that harbors the GOF mutation (Fig. 1A). To characterize this mouse model, we first generated constitutive GOF *Piezo2* mice (GOF^const.^) by breeding mice homozygous for the GOF allele into *Cmv*^Cre^ driver that expresses Cre recombinase ubiquitously (11). To validate our GOF model at the cellular level, we used whole-cell patch clamp to record PIEZO2-dependent mechanically activated (MA) currents from dorsal root ganglion (DRG) neurons of homozygous GOF *Piezo2* mice and wild type littermates (Fig. S1A). We initially tested homozygous instead of heterozygous mice to avoid diluting the effects of the GOF allele by a wildtype allele. Although the majority of homozygotes died perinatally for unknown reasons, some survived and we were able to obtain DRGs from viable mice for recording. DRG neurons from homozygous GOF^const.^ mice showed a significantly slower inactivation and larger steady state currents compared to wild type (Fig. 1B and C). As expected, there was no detectable difference in I_max_ and apparent threshold between wild type and GOF neurons (Fig. S1B). We next set out to compare MA DRG currents of heterozygous mice to wild type to reflect the human condition more closely. To enrich the recorded population for PIEZO2-dependent rapidly inactivating neurons, we crossed mice carrying the GOF *Piezo2* allele into *Pvalb^Cre^/Ai9*, which express Cre recombinase and Ai9 tdTomato reporter protein in proprioceptive neurons (7, 12). DRG neurons from these mice were isolated and tdTomato+ cells were subjected to whole-cell patch clamp recordings (Fig. 1D and Fig. S1C). As expected, we observed robust differences in the kinetics of MA currents between heterozygous GOF and wild type DRGs (Fig. 1D and Fig. S1C). These results validate that the transgenic mice express a functional GOF PIEZO2 channel with similar kinetics of inactivation as human GOF mutations analyzed in heterologous system (5). They also demonstrate that heterozygosity of GOF PIEZO2 is sufficient to increase DRG mechanosensitivity *in vivo*.

**Figure 1.**
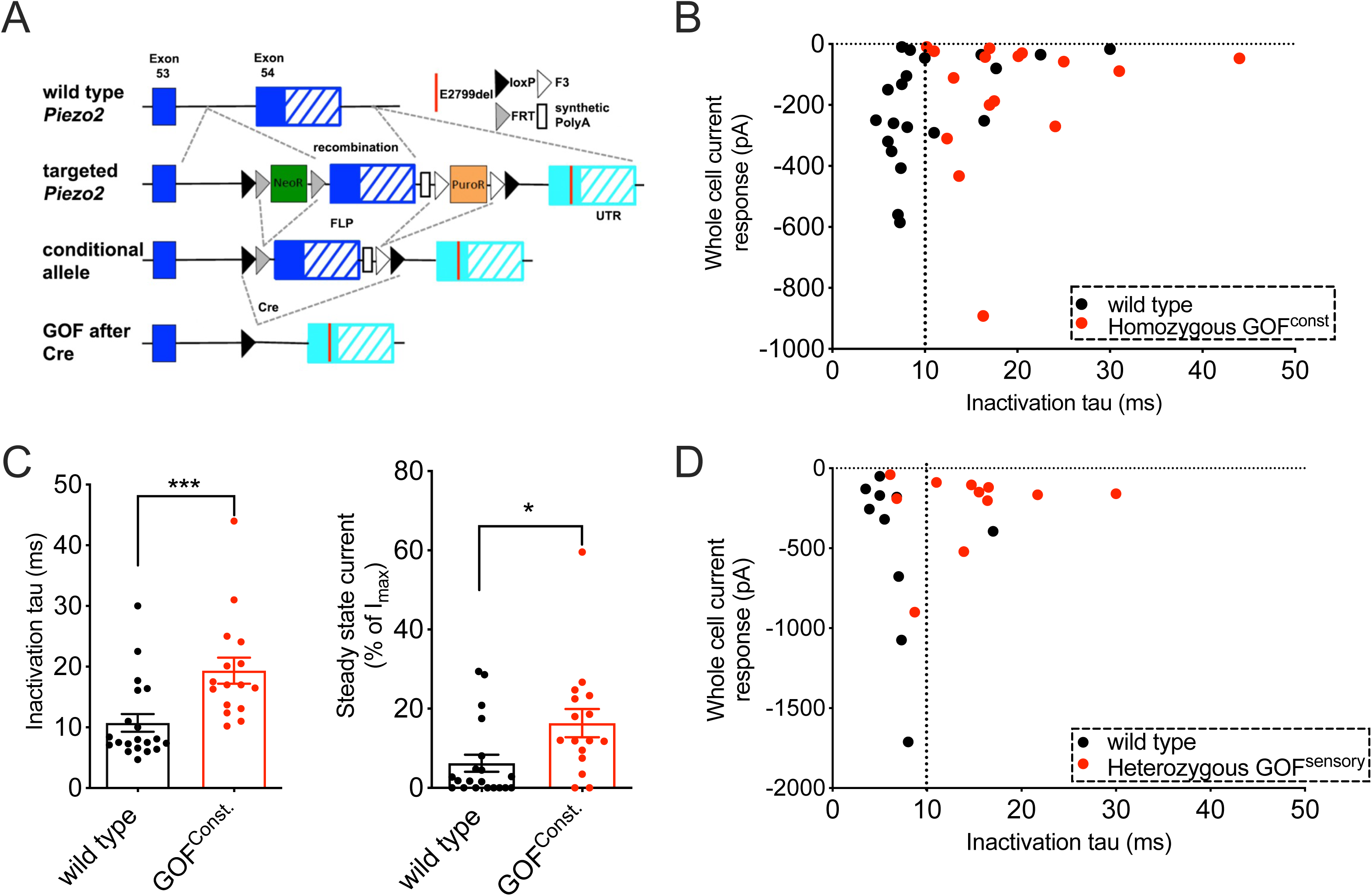
Human equivalent gain-of-function (GOF) PIEZO2 increases mechanosensitivity of sensory neurons in mice. (A) Schematic description of strategy for generating conditional GOF *Piezo2* mice. Briefly, in cells that express Cre recombinase, the wild type exon (blue) will be replaced by the modified exon (Cyan) that harbors the E2799del GOF point mutation. (B) Mechanically activated currents and inactivation time constant (tau) in DRG sensory neurons from wild type (black) and constitutively homozygous GOF *Piezo2* mice (red). 87% WT and 70% GOF neurons were responsive to the stimulus. (C) Bar graphs representing inactivation time constant (ms) and steady state current (% of Imax value) in DRG sensory neurons. (D) Mechanically activated currents and inactivation time constant in Tdtomato+ proprioceptive sensory neurons from *Pvalb^Cre^*/*Ai9* (wild type, black) and GOF *Piezo2; Pvalb^Cre^/Ai9* mice (red). *p < 0.05, **p < 0.01, and ***p < 0.001 (Student’s t -test). Each data point represents a single DRG neuron.

To test whether GOF PIEZO2 causes DA5-like phenotypes in mice, and to study the underlying mechanisms, we analyzed the constitutive GOF^const.^ mice. Because DA5 patients have one allele of GOF *PIEZO2*, we used heterozygous GOF^const.^ mice as the model. We observed that hindlimbs of these mice had contractures compared to wild type littermates whose hindlimbs had well-extended digits (Fig. 2A). Micro-computed tomography (CT) images showed that the angles between major hindlimb joints were smaller in GOF^const.^ mice than in wild type (Fig. 2A). Specifically, we measured the angle between the phalange and metacarpal, the two major bones of hindlimbs. We found that phalange-metacarpal angles in hindlimbs of GOF^const.^ mice were significantly decreased compared to wild type mice (Fig. 2B). In addition to joint abnormalities, DA5 patients also have weak or absent tendon reflexes suggesting that defective tendon development might be associated with joint contractures (5). To evaluate tendon phenotypes, we dissected the intact tendons from hindlimbs (Fig. 2C), and we observed that the average length of tendons was significantly reduced in GOF^const.^ mice compared to wild type (Fig. 2D). It is therefore possible that the DA5-like contractures are caused by shortened tendons. Our analysis shows that GOF *Piezo2* mice have anatomical defects similar to those observed in DA5 patients.

**Figure 2.**
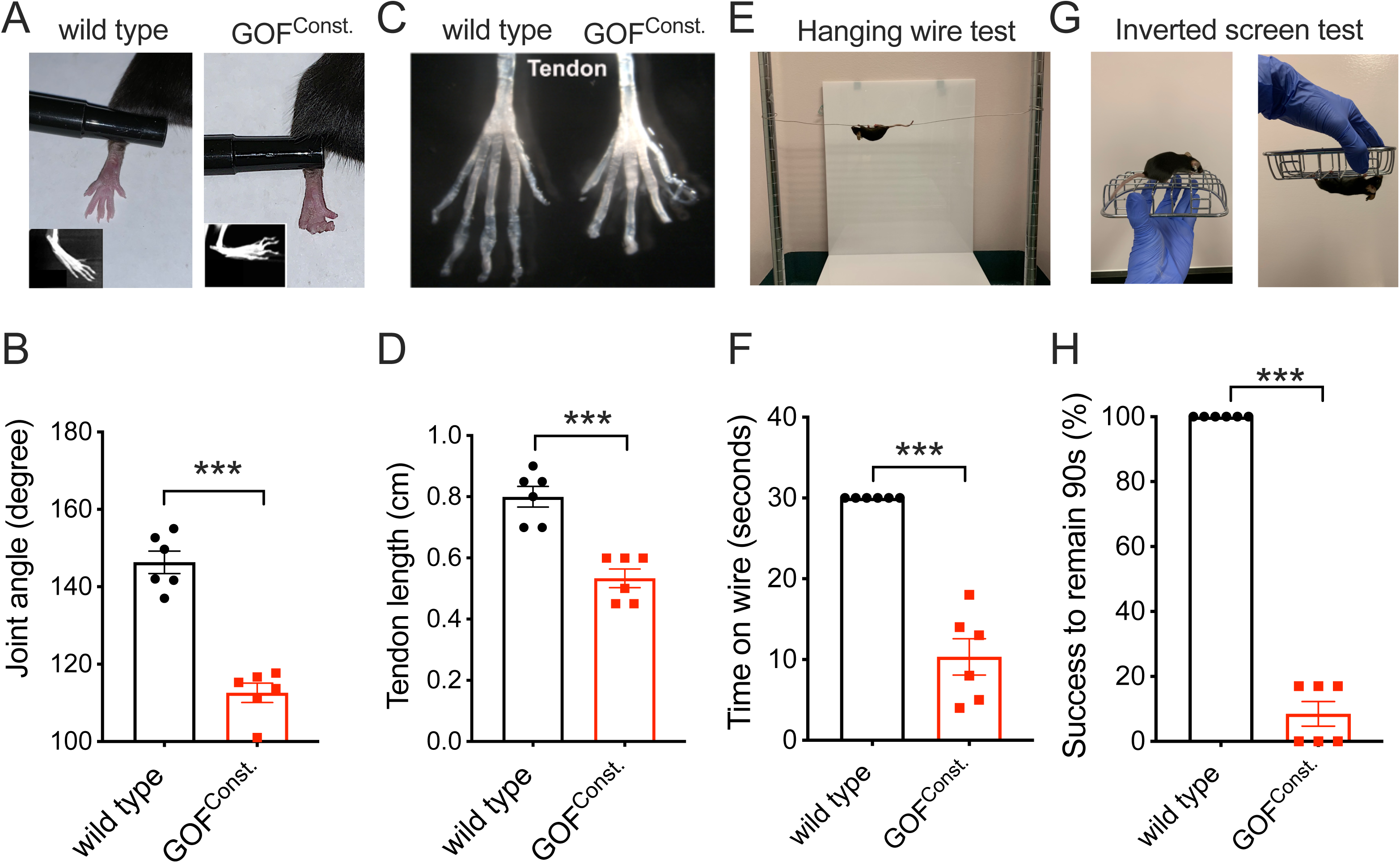
GOF *Piezo2* mice develop anatomical defects and compromised limb function. (A) Joint morphology in the hindlimb of wild type and constitutive heterozygous GOF *Piezo2* mice. (B) Quantification of phalange-metacarpal joint in mice. (C) Intact tendon dissected from wild type and constitutive heterozygous GOF *Piezo2* mice. (D) Quantification of tendon length. (E) Image representing hanging wire test. (F) Time (sec) that animals remained on the metal wire in (E), with 30 sec as cutoff. (G) Image representing inverted screen test. (H) Quantification for (G) as percentage of animals successfully remaining on the rotating screen for 90 seconds. *p < 0.05, **p < 0.01, and ***p < 0.001 (Student’s t-test). Each data point represents a single animal.

Anatomical deformities cause limited joint movements and affect motor functions in distal arthrogryposis patients. To test whether GOF PIEZO2 impacts limb functionality in mice, we performed two behavioral tests. First, we used a hanging-wire assay to evaluate limb functionality in both wild type and GOF^const.^ mice (13). The test is based on the latency of a mouse to fall off from a metal wire (Fig. 2E). We found that wild type mice were able to remain on the wire for at least 30 seconds, whereas GOF^const.^ mice fell off significantly earlier (Fig. 2F). Second, we performed the inverted screen test (14), which is another method of evaluating limb functionality in mice. This assay measures the time until a mouse falls off a square screen after it is rotated upside down (Fig. 2G and method). We tested each mouse on this assay six times and calculated the success rate for remaining on the rotating screen for at least 90 seconds. All wild type mice were able to hold on for at least 90 seconds. In contrast, the average success rate for GOF^const.^ mice was less than 10% (Fig. 2H). Thus, these results suggest that limb functionality was compromised in GOF^const.^ mice and this is consistent with the severe limb anatomical defects in these mice.

To investigate which cell types are involved in the DA5 etiology, we expressed GOF *Piezo2* allele in various tissues. Although PIEZO2 is not required in muscles and other connective tissues during development (15), it is still possible that ectopic activity of moderate levels of this ion channel could cause musculoskeletal deficits. To test this, we induced GOF PIEZO2 expression in skeletal muscle or all mesenchymal cells including cartilages and tendons (in addition to muscles) by crossing mice carrying the conditional GOF *Piezo2* allele with *MCK*^Cre^ (16), for skeletal muscle and *Prrx1*^Cre^ (17), for mesenchyme, respectively. These GOF *Piezo2* mice had normal phalange-metacarpal angles and tendon lengths in their hindlimbs compared to wild type littermate controls (Fig. 3A and B), suggesting that overactive PIEZO2 in mesenchyme-derived connective tissues is unlikely to cause joint contractures in mice. Consistently, these mice performed comparably to wild type mice on both the hanging wire and inverted screen behavioral assays (Fig. 3C and D). Remarkably, we found that expression of GOF PIEZO2 in proprioceptive sensory neurons (via *Pvalb^Cre^*mice) that mainly innervate muscle spindles and tendon organs (7) was sufficient to cause joint contractures and shortened tendons with similar severity observed in constitutive GOF *Piezo2* mice (Fig. 3A and B). In addition, we observed that these sensory neuron-specific GOF *Piezo2* mice had poor performance on both the hanging wire and the inverted screen behavioral assays (Fig. 3C and D), comparable to constitutive GOF *Piezo2* mice. These data suggest that overactive PIEZO2 in sensory neurons, but not in muscle, bone, cartilage or tendon, is sufficient to induce DA5-like phenotypes *in vivo*.

**Figure 3.**
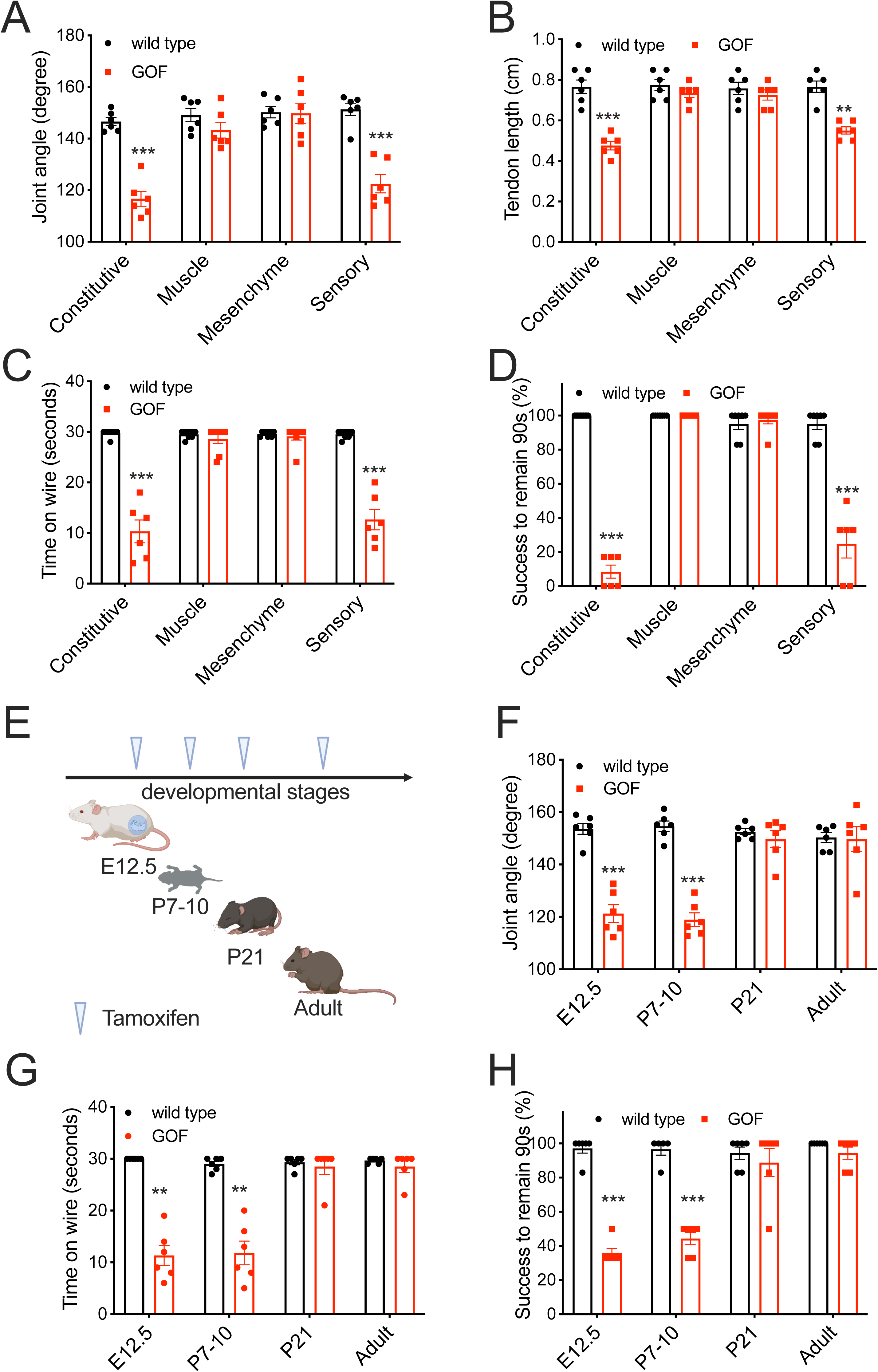
GOF PIEZO2 in sensory neurons is sufficient to cause defects in limb anatomy and functions during a critical developmental stage. (A) Phalange-metacarpal joint angle of the hindlimbs in wild type, constitutive and tissue specific GOF *Piezo2* mice (black: wild type; red: GOF *Piezo2*). (B) Tendon length of hindlimbs in mice. (C) Quantification of hanging wire test results for mice. (D) Quantification of inverted screen test results for mice. (E) Tamoxifen induction of GOF PIEZO2 in proprioceptive neurons by *Advillin^Cre-ERT2^*at various developmental stages. (F) Phalange-metacarpal joint angle of the hindlimbs in wild type and GOF *Piezo2*; *Advillin^Cre-ERT2^* mice injected with tamoxifen at different developmental stages. (G) Quantification of hanging wire test results for wild type and GOF *Piezo2; Advillin^Cre-ERT2^*mice. (H) Quantification of inverted screen test for wild type and GOF *Piezo2; Advillin^Cre-ERT2^* mice. *p < 0.05, **p < 0.01, and ***p < 0.001 (One-way ANOVA followed by Tukey’s multiple comparison). Each data point represents a single animal.

Since DA is a developmental disorder presumably originating during fetal stages (2), we used our mouse model to determine the timing of PIEZO2 activity in sensory neurons that affects musculoskeletal development. We used an inducible sensory neuron-specific GOF *Piezo2* mouse by crossing the GOF allele into *Advillin*^Cre-ERT2^ driver mice (18). Advillin, an actin binding protein, is expressed in most DRG neurons, but not central nervous system, starting in early embryogenesis (19). We induced overactive PIEZO2 expression in sensory neurons at various developmental stages by tamoxifen injection (Fig. 3E). When we induced the expression of GOF *Piezo2* allele at embryonic day 12.5 or at postnatal day 7-10 (P7-10), these mice showed significantly decreased phalange-metacarpal joint angles (Fig. 3F), suggesting that the phenotype is mainly dependent on expression of PIEZO2 sometime after P7. However, induction of GOF PIEZO2 expression in sensory neurons after P21 or during adulthood (> 1month old) did not cause joint defects in mice (Fig. 3F). Consistent with anatomical phenotypes, mice expressing GOF PIEZO2 in sensory neurons during embryonic (E12.5) and early postnatal stage (P7-10), but not after early adulthood (>P21), performed poorly on both hanging wire and inverted screen tests (Fig. 3G and H). These results confirm our earlier findings using *Pvalb*^Cre^ that overactive PIEZO2 in sensory neurons causes DA5-like defects (Fig. 2). In addition, these experiments suggest a critical postnatal developmental period between days 7 and 21 during which excessive mechanotransduction in somatosensory neurons is the primary cause of the limb malformation that we observe.

To ensure that the anatomical deficits we observed are not due to an indirect consequence of compromised proprioceptive neuronal development we examined muscle spindles, a major subtype of proprioceptive nerve endings anatomically. We did not observe overt anomalies in these proprioceptive endings that could potentially be a primary cause of tendon malformations in these mice (Fig. S2).

To further test whether anatomical defects of GOF *Piezo2* mice are attributed to hyperactivation of neurons, we performed intra-muscular injections of Botulinum neurotoxin type A (commonly known as Botox). This neurotoxin is known to inhibit neurotransmitter and peptide secretion in a variety of neuronal types by cleaving SNAP25, a SNARE protein essential for vesicular exocytosis (20). We injected Botox into the hindlimb of mice at P7-10, at the beginning of the critical period (Fig. 4A). The contralateral hindlimb received vehicle injection as an internal control. Four weeks after a single dose of injection, we observed that Botox did not affect joint morphology in wild type mice (Fig. 4B). Remarkably, however, it rescued joint defects in GOF *Piezo2* hindlimbs compared to vehicle controls (Fig. 4B). This result is consistent with neuronal activity to be responsible for the joint defects we observe.

**Figure 4.**
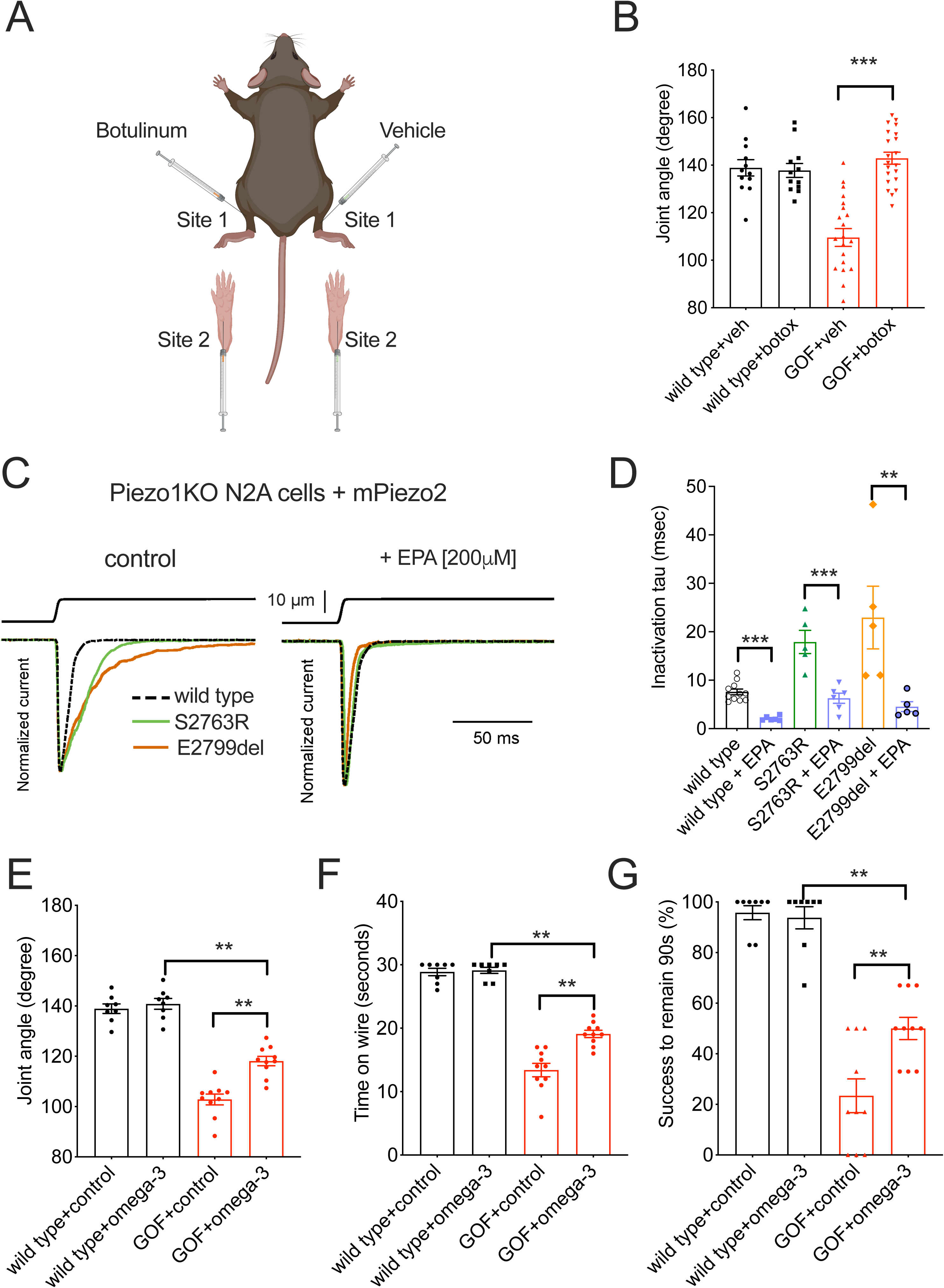
Local injection of Botulinum toxin and dietary intake of EPA rescue limb defects in GOF *Piezo2* mice. (A) Carton describing Botulinum toxin injection into two sites of a single hindlimb in mice of P7-10. The other hindlimb of the same animal injected with vehicle served as control. (B) Phalange-metacarpal joint angle of the hindlimbs in young adult wild type and GOF *Piezo2*; *Pvalb^Cre^* mice that received Botulinum toxin at P7-10. (C) Whole cell poke induced MA currents in N2A*Piezo1*-/- cells heterologously expressing mouse PIEZO2 wild type (stippled) and GOF mutants S2763R (green) and E2799del (brown). (D) Quantification of the inactivation time constant results shown in C. (E) Phalange-metacarpal joint angle of the hindlimbs in young adult wild type and GOF *Piezo2*; *Pvalb^Cre^*mice that received EPA-enriched diet. Sunflower-oil -based food, which has a similar level of fat content and energy, used as a control. (F) Quantification of hanging wire test results in mice fed with EPA-enriched diet. (G) Quantification of inverted screen test results in mice fed with EPA-enriched diet . *p < 0.05, **p < 0.01, and ***p < 0.001 (Two-way ANOVA followed by Tukey honestly significance test). Each data point represents a single hindlimb in B; a single neuron in D; and a single animal in E – G.

Our data suggests that any manipulation that would decrease overall PIEZO2 activity during a critical development time could potentially rescue the DA5-like phenotype. The μ-3 polyunsaturated fatty acid (PUFA) eicosapentaenoic acid (EPA), a membrane lipid component, significantly decreases the inactivation time of wild type and GOF PIEZO1 channels, a close relative of PIEZO2 (21, 22). Because EPA is commonly found in fish and widely used as a dietary supplement, we wanted to test whether feeding mice a diet enriched in μ-3 PUFAs has therapeutic potential for treating DA5-like defects via modulating PIEZO2 activity.

We first showed that overnight incubation with EPA (200 µM) significantly reduced wild type as well as GOF mutant PIEZO2 inactivation time constant in a heterologous expression system (Fig. 4C and D). This suggests that EPA has the potential to modulate disease causing mutations in PIEZO2. To test the potential of EPA *in vivo*, female mice were fed a diet enriched in EPA during pregnancy and nursing and the sensory neurons of their offspring were assessed for PIEZO2-dependent mechanosensitivity (Fig. S3A). We found that EPA -enriched diet intake significantly reduced inactivation time constant of mechanically stimulated currents in sensory neurons but had no effect on the current intensity and apparent threshold (Fig. S3A). Next, we determined the effect of this diet on the anatomical defects in GOF *Piezo2* mice. Liquid chromatography-mass spectrometry (LC-MS) showed that the feeding protocol successfully incorporated EPA into the plasma membrane of sensory neurons that carry GOF *Piezo2* mutation (Fig. S3B). Remarkably, dietary EPA partially prevented joint malformations (Fig. 4E) and improved the performances on the hanging wire and inverted screen tests (Fig. 4F and G) in GOF *Piezo2* mice. These results are consistent with the hypothesis that overactive PIEZO2 signaling causes the DA5-like phenotype.

Overall, our study has identified a critical period during development when relatively small excess PIEZO2 activity in these neurons can cause severe joint contractures in distal arthrogryposis. Thus, precise spatiotemporal control of mechanosensation in the peripheral nervous system is crucial for musculoskeletal development. Previous work has shown that loss of PIEZO2 in proprioceptive neurons causes defective spinal alignment in mice (15). While a requirement of proprioceptive feedback for proper spinal alignment is easy to envision, the adverse effect of excessive mechanotransduction in proprioceptive neurons on limb anatomy is unexpected. Mechanosensory activation of proprioceptive nerve endings causes an efferent, Ca^2+^- dependent signaling within the proprioceptive endings culminating in exocytosis (23, 24). It also causes afferent signaling and firing of proprioceptive neurons that will activate downstream motor neurons innervating muscles (25). Further work is needed to determine the contribution of the efferent and afferent signaling pathways to anatomical deficits in GOF *Piezo2* mice.

Importantly, we present the first proof-of-concept that Botox injection or dietary treatment can counteract the effect of overactive PIEZO2 function to evade DA-like phenotypes in mice when applied during a critical period. These approaches might have clinical applications for human disorders caused by GOF PIEZO2 mutations. Beyond this, our findings call attention to mechanotransduction of the sensory system when diagnosing and treating other musculoskeletal disorders.

## Acknowledgements

We thank Anton Maximov for discussion and for critical reading of the manuscript. This work was supported by the following grants: NIH R35 NS105067 and R01 DE022358 to A.P. A.P. is an investigator of the Howard Hughes Medical Institute. The authors declare no competing financial interests.

## Materials and Methods

### Animals

All animal procedures were approved by the Institutional Animal Care and Use Committees of The Scripps Research Institute (California) and the University of Tennessee Health Science Center. Mice carrying the conditional gain-of-function (GOF) *Piezo2* allele were generated and maintained on C57BL/6 background. All animals were backcrossed at least 10 generations into C57BL/6. Constitutive GOF *Piezo2* (GOF^const.^) mice were generated by breeding mice with conditional GOF allele to *Cmv^Cre^* driver (The Jackson Laboratory, stock #006054). Tissue specific GOF *Piezo2* mice were generated by breeding conditional GOF mice into *MCK^Cre^* (The Jackson Laboratory, stock#006475), *Prrx1^Cre^*(The Jackson Laboratory, stock#005584), *Pvalb^Cre^* (The Jackson Laboratory, stock#017320) and *Advillin*^Cre-*ERT2*^ (The Jackson Laboratory, stock#026516) driver mice, respectively. Reporter mouse Ai9 was from The Jackson Laboratory, stock#007909. Mice were housed in a temperature-controlled (22-24°C) room that maintains a 12hr light/dark cycle. All tissue harvests and behavioral assays were performed on mice between 1.5 to 6 months old. Pharmacological treatments were done on mice at different ages as specified (see below for details).

### Electrophysiology and mechanical stimulation

#### DRG neurons

Mechanically activated currents from DRG neurons 1-3 days after culturing were recorded in whole cell patch clamp mode using a MultiClamp700A amplifier and DigiData1550 (Molecular Devices) and stored directly and digitized online using pClamp software (version 10.7). Currents were sampled at 20 kHz and filtered at 2 kHz. Recording electrodes had a resistance of 3 to 7 Megohms when filled with gluconate-based low-chloride intracellular solution: 125 mM K-gluconate, 7 mM KCl, 1 mM CaCl_2_, 1 mM MgCl_2_, 10 mM HEPES (pH with KOH), 1 mM tetra-K BAPTA [1,2- bis(2-aminophenoxy)ethane-N,N,N′,N′-tetraacetic acid], 4 mM Mg-ATP (adenosine triphosphate), and 0.5 mM Na-GTP (guanosine triphosphate). Extracellular bath solutions used: 133mM NaCl, 3mM KCl, 2.5mM CaCl_2_, 1mM MgCl_2_, 10mM HEPES (pH7.4), and 10mM glucose.

Mechanical stimulation was achieved using a fire-polished glass pipette (tip diameter, 3 to 4 µm) positioned at an angle of 80° relative to the cell being recorded. Displacement of the probe toward the cell was driven by Clampex-controlled piezoelectric crystal microstage (E625 LVPZT Controller/Amplifier; Physik Instrumente). The probe had a velocity of 1 µm msec−1 during the ramp phase of the command for forward movement, and the stimulus was applied for a duration of 125 msec. For each cell, a series of mechanical steps in 1-µm increments was applied every 10 sec.

#### Heterologous expression in N2A^Piezo1-/-^ cells

For whole-cell recordings of mechano-activated currents in Piezo1-/- N2A cells and mouse pups DRG neurons in EPA -related experiments, the bath solution contained 140 mM NaCl, 6 mM KCl, 2 mM CaCl2, 1 mM MgCl2, 10 mM glucose, and 10 mM HEPES (pH 7.4), while the pipette solution contained 140 mM CsCl, 5 mM EGTA, 1 mM CaCl2, 1 mM MgCl2, and 10 mM HEPES (pH 7.2). Pipettes were made of borosilicate glass (Sutter Instruments) and fire-polished to a resistance between 3 and 5 MΩ before use. Mechanical stimulation was performed using the voltage-clamp (constant -60 mV) configuration. Recordings were sampled at 100 kHz and low pass filtered at 10 kHz using a MultiClamp 700 B amplifier and Clampex (Molecular Devices, LLC). Leak currents before mechanical stimulations were subtracted offline from the current traces, and data were digitally filtered at 2 kHz with ClampFit (Molecular Devices, LLC).

Recordings with leak currents > 200 pA, with access resistance >10 MΩ, and cells with giga-seals that did not withstand at least five consecutive steps of mechanical stimulation were excluded from analyses.

#### Other buffers

CsCl-based intracellular solution: 133 mM CsCl, 1 mM CaCl2, 1 mM MgCl2, 10 mM HEPES (pH with CsOH), 5 mM EGTA, 4 mM Mg-ATP (adenosine triphosphate), and 0.5 Na-GTP (guanosine triphosphate). Extracellular bath solution was composed of 133 mM NaCl, 3 mM KCl, 2.5 mM CaCl2, 1 mM MgCl2, 10 mM HEPES (pH 7.3 with NaOH), and 10 mM glucose.

### N2A cell transfections

Neuro2A cells were co-transfected with 250-500 ng*mL^-1^ of mouse Piezo2 cDNA (wild type and S2691R and E2727del GOF mutants) and 50 ng/mL GFP-pMO, using Lipofectamine 2000 (Thermo Fisher Scientific) according to the manufacturer’s instructions. Fatty acids were supplemented 18-24 hours before recording.

### Liquid chromatography-mass spectrometry

DRG tissues were frozen in liquid nitrogen immediately after dissection from GOF *Piezo2* mice fed with standard, sunflower, or EPA-enriched diets. Total and free fatty acids were extracted and quantified at the Lipidomics Core Facility at Wayne State University. Membrane (i.e., esterified) fatty acids were determined by subtracting free from total fatty acids and normalized by protein content

### Anatomical analyses

Wild type and GOF mice littermates between 1.5 to 6 months old were euthanized by isoflurane. Distal part (feet) of hindlimbs from the mice was severed, with skins and calcaneus (heel bone) intact, followed by fixation in 4% PFA for one week at 4°C. The angle between phalange and metacarpal was measured for the middle three digits. The data was shown as the average for each mouse.

For tendon analysis, the intact flexor tendon was detached and dissected out from the bones and muscles of mouse hindlimb. The tendon length was measured as the distance from the tip of third (longest) flexor tendon to the attachment site to the flexor digitorum profundus.

### Immunohistochemistry

Adult mice were perfused with phosphate-buffered saline (PBS) and 4% paraformaldehyde (PFA) followed by isolation of the skeletal muscle epitrochleoanconeus (ETA) from the upper forelimbs. Whole muscles were post-fixed in 4% PFA overnight in a rotating incubator in 4°C. Muscles were washed in 30mL PBS for 2h followed by 30 mL PBS with 0.3% TritonX-100 (PBST) for 6h. Whole muscles were blocked in 1% bovine serum albumin (BSA, wt/vol), 5% normal goat serum (NGS, vol/vol), and 0.3% PBST for 1h. Primary antibodies were incubated in 1% BSA, 5% NGS, and 0.3% PBST for 3 days at 4°C on rotating incubator (NFH, 1:500, Abcam, ab4680; Tubulin β-III, 1:200, Biolegend, 801201). Whole muscles were washed in 0.3% PBST for 5 hours then incubated in secondary antibodies for 48h at 4°C on rotating incubator (AlexaFluor 568, 1:200, Invitrogen, A11041; AlexaFluor 647, 1:200, Invitrogen, A21236). Whole muscles were dehydrated in a series of 25%, 50%, and 75% MeOH in PBS for one hour each on shaking incubator at room temperature, protected from light. Whole muscles were dehydrated in 100% MeOH overnight on shaking incubator at room temperature. Muscles were incubated in 1-part benzyl alcohol to 2-part benzyl benzoate for 1 hour then whole mounted for imaging.

### Behavioral tests

#### Hanging wire assay

A 60cm long and 2mm thick metal wire was secured to two vertical poles. The wire was tightly attached to the poles to avoid displacement and vibration while the mice were being assayed on the wire. Also, the wire was about 40cm above a layer of soft bedding. The suspension time for each mouse was measured in three trials with one minute recovery period between trials, and the average was calculated. The time until the animal completely fell off from its grasp was recorded. Because all wild type mice were able to remain on the wire for at least 30 seconds while GOF mice had significantly shorter suspension time, 30-second was used as a cutoff.

#### Inverted screen test

Mice were placed individually on the top of a metal square (around 8cm x 8cm) grid screen. The screen was then rotated 180° and the mice were on the bottom of the screen. The suspension time on the screen (either remaining on the bottom or climbing up to the top side) was recorded for each mouse in six trials, with one minute recovery period between trials. The percentage of successful trials with at least 90-second suspension time was calculated for each mouse.

### Pharmacological experiments

Botulinum toxin A (Botox, 100 units) was purchased from Allergan (Irvine, CA) and diluted by water to a working solution of 5U/ml. Wild type and GOF *Piezo2*; *Pvalb^Cre^* mice at P7-10 were briefly immobilized on ice immediately before drug injection. Botox was injected into two sites of the hindlimb: one dose (0.5U kg^-1^) via intramuscular injection into distal part of gastrocnemius and another dose (0.5U kg^-1^) via intraplantar injection into the paws. For each animal, one side of hindlimb received drug injection while the contralateral side was injected with vehicle as an internal control.

### Tamoxifen injection

Tamoxifen was administered as previously published (6). Briefly, 150mg of tamoxifen (Sigma) was dissolved in 10ml of 100% corn oil (at 50°C) freshly daily before use. Dissolved tamoxifen solution was injected intraperitoneally (at room temperature) into wild type and GOF *Piezo2*; *Advillin^Cre-ERT2^* mice (littermates) (see main text) at 150mg kg^-1^ for five consecutive days. Each mouse was weighted before injection to normalize for differences in body weight. Anatomical analyses and behavioral assays were performed between 4 to 6 weeks after tamoxifen injection.

### Diet intervention

Breeding mice were fed with standard, menhaden (high EPA content, Ref), or sunflower (a control diet with similar caloric content) oil-enriched diets. For electrophysiology experiments, pregnant females, when reaching late pregnancy (2-3 days before birth), were relocated to a new cage and fed with regular rodent diet until their pups were 9 days old. The reason for this relocation was to reduce cannibalism observed when newborns and parents were in the same EPA-enriched diet feeding cage. DRGs from P9 pups were dissected, cultured, and for mechanically activated (MA) currents. For analyses on mouse anatomy and behavior, parenting mice and newborn pups were housed in the same EPA-enriched diet feeding cage with a different bedding material. Cannibalism was reduced by this change and relocation was not necessary (i.e. these animals were fed EPA-enriched diet throughout their life). Anatomical and behavioral assays were performed when pups were 5 weeks old.

### Statistical analysis

Statistical analyses were performed and significance was computed using GraphPad Prism 6. Student’s t tests were used when comparison was made between two groups. Unpaired *t* test with Welch correction was used when comparing three groups. One-way and two-way ANOVA with multiple comparison tests were performed when comparing two groups, at multiple time or conditions. When specific tests have been made, it is stated in the figure legend. P values (ɑ = 0.05, two sided) are indicated in figure legends.

## Supplementary figure legends

**Figure S1.**
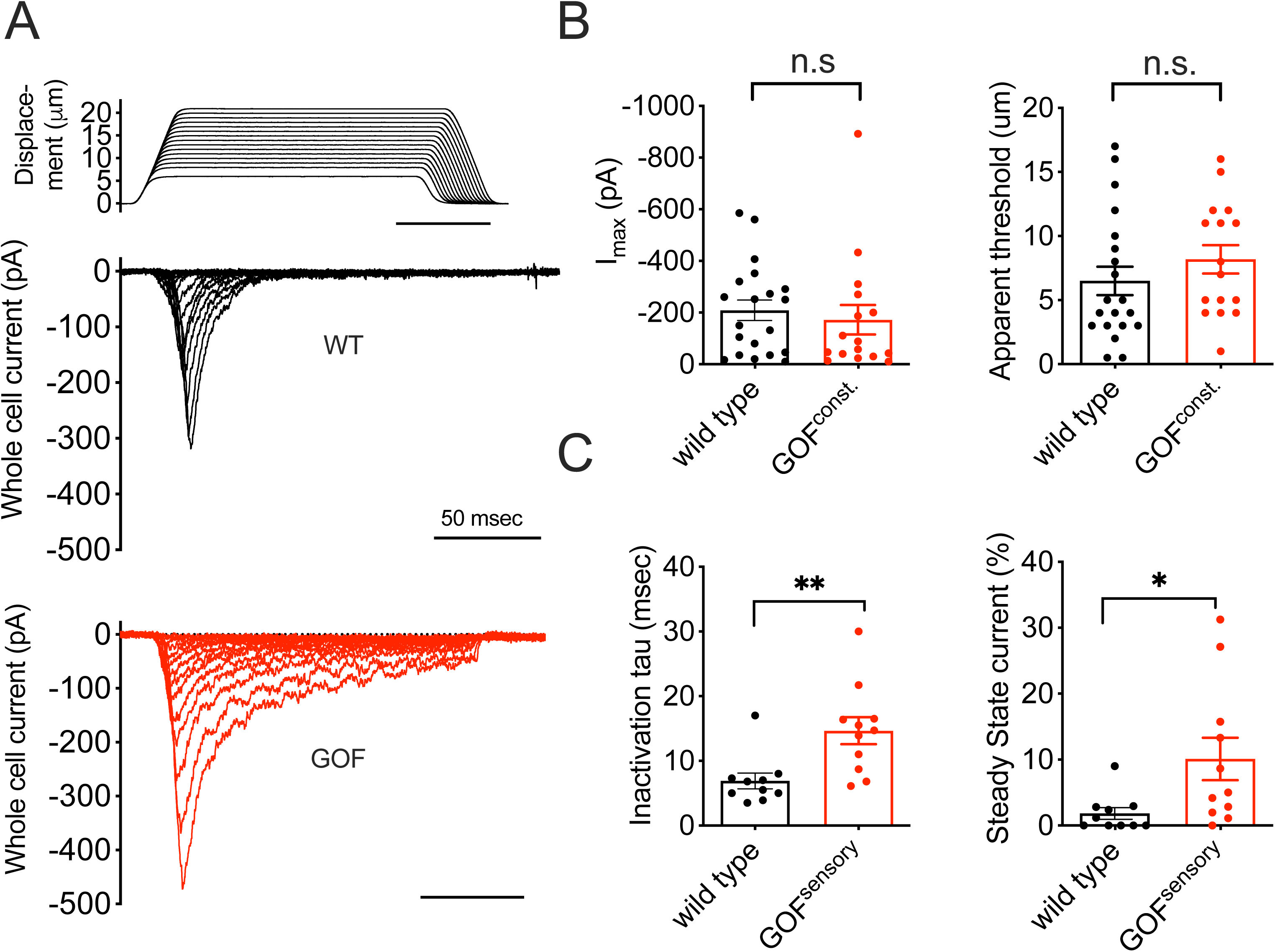
(A) Representative traces of mechanically activated (MA) currents (whole - cell patch clamp recording) for wild type (WT, black) and homozygous constitutive GOF *Piezo2* DRG (GOF^const.^, red) sensory neurons. (B) Quantification for I_max_ and apparent threshold of wild type (WT) and homozygous GOF^const.^ DRG neurons. (C) Inactivation time constant (tau) and steady state current of MA currents from wild type (WT) and heterozygous GOF *Piezo2*; *Pvalb^Cre^*(GOF^sensory^) DRG neurons.

**Figure S2.**
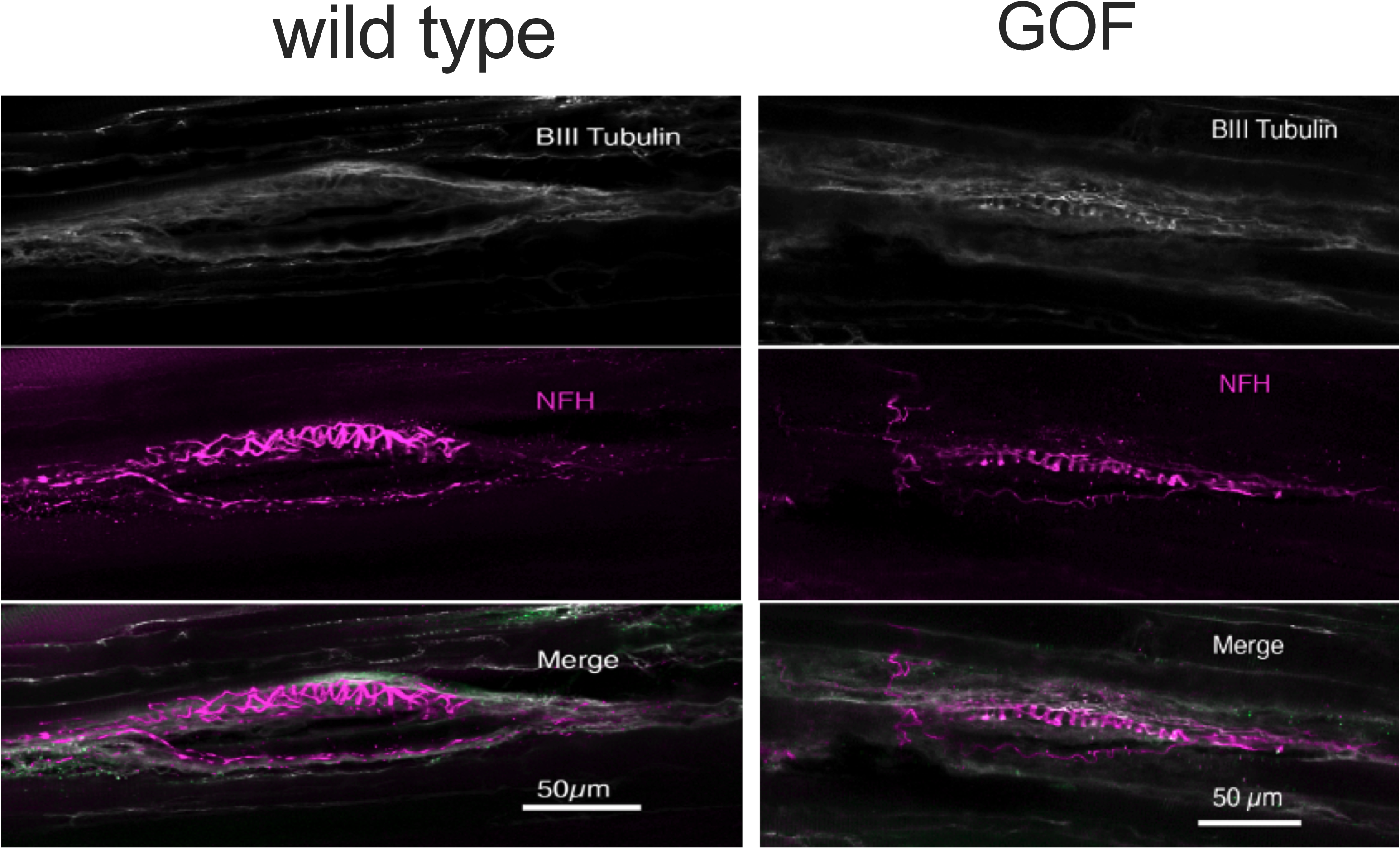
Morphology of proprioceptors in the skeletal muscles of wild type (WT) and GOF *Piezo2*; *Pvalb^Cre^* mice. Neuronal makers Beta-tubulin III (BIII tubulin; white) and neurofilament heavy chain (NFH; purple) immunohistochemistry showing normal spindle morphology (merged image). Representative images from three pairs of wild type and GOF mice. Scale bar: 50 μm.

**Figure S3.**
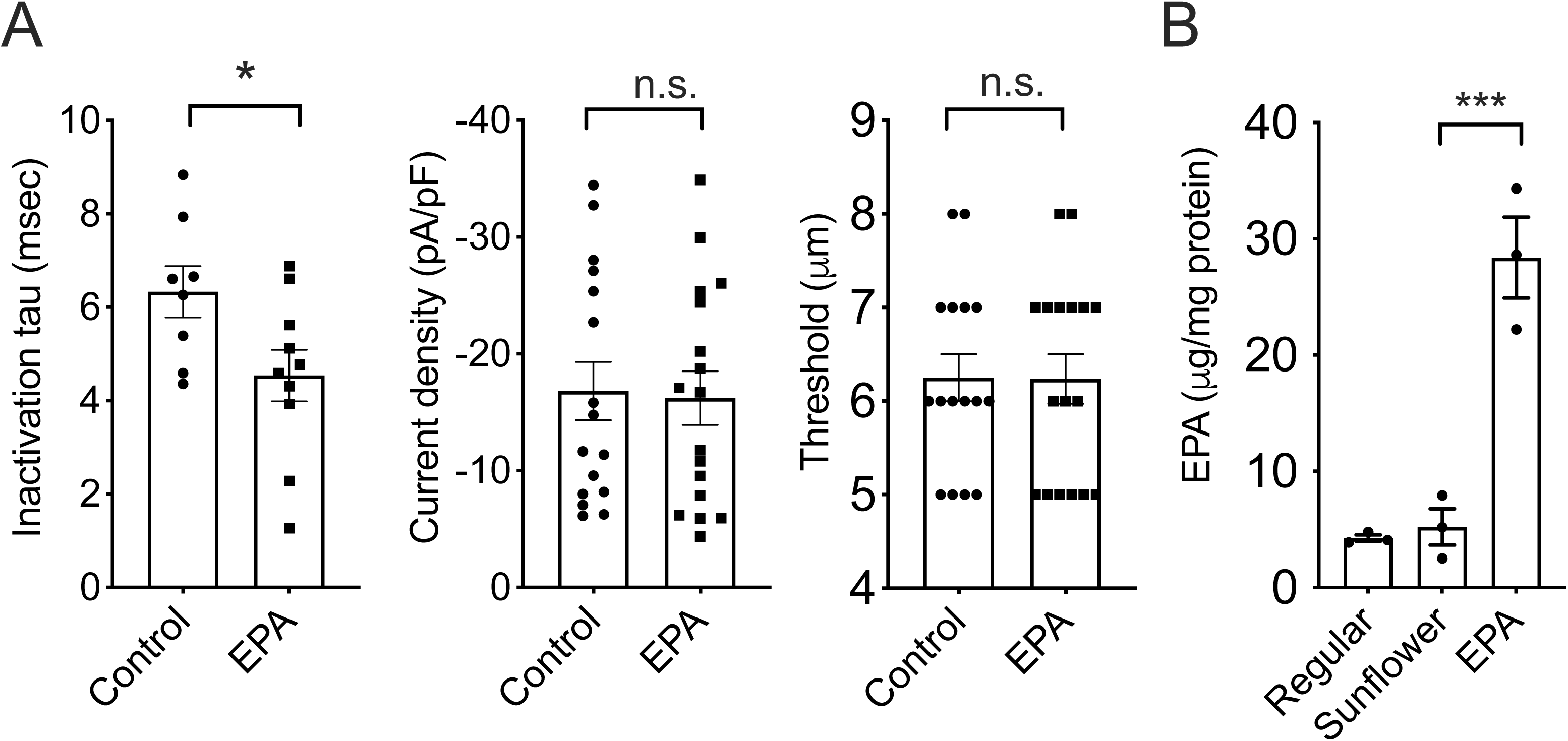
(A) Whole cell recordings from wild type mice. EPA decreases the inactivation time constant of rapidly inactivating PIEZO2-dependent currents in DRG neurons (left) but has no effects on MA current magnitudes (I_max_, middle) and apparent thresholds (right). (B) EPA membrane content of DRG from mice fed with standard, sunflower, or menhaden oil (μ-3 EPA) enriched diets, as determined by LC-MS (Unpaired *t* test with Welch correction).

